# Inhibition of SQSTM1/p62 oligomerization and Keap1 sequestration by the Cullin-3 adaptor SHKBP1

**DOI:** 10.1101/2025.01.21.634088

**Authors:** Lin Luan, Xiaofu Cao, Jeremy M. Baskin

## Abstract

SQSTM1/p62 is a master regulator of the autophagic and ubiquitination pathways of protein degradation and the antioxidant response. p62 functions in these pathways via reversible assembly and sequestration of additional factors into cytoplasmic phase-separated structures termed p62 bodies. The physiological roles of p62 in these various pathways depends on numerous mechanisms for regulating p62 body formation and dynamics that are incompletely understood. Here, we identify a new mechanism for regulation of p62 oligomerization and incorporation into p62 bodies by SHKBP1, a Cullin-3 E3 ubiquitin ligase adaptor, that is independent of its potential functions in ubiquitination. We map a SHKBP1–p62 protein-protein interaction outside of p62 bodies that limits p62 assembly into p62 bodies and affects the antioxidant response by preventing sequestration and degradation of Keap1. These studies provide a non-ubiquitination-based mechanism for an E3 ligase adaptor in regulating p62 phase separation and cellular responses to oxidative stress.

## INTRODUCTION

Protein degradation and quality control are fundamental to cellular health, with autophagy and the ubiquitin-proteasome system (UPS) serving as two major regulatory mechanisms (Pohl and Dikic, 2019). The molecular integration of these pathways is mediated by autophagy receptors, which simultaneously bind ubiquitinated proteins and autophagy-specific ubiquitin-like modifiers such as LC3/GABARAP (Kirkin et al., 2009). The best studied autophagy receptor is p62/SQSTM1, whose dysfunction is closely linked to multiple neurodegenerative diseases (Ma et al., 2019; Rubinsztein, 2006) and cancers (Moscat et al., 2016). p62 participates in two critical cellular processes: the clearance of ubiquitinated proteins via autophagy (Pankiv et al., 2007) and the activation of antioxidant responses through the Keap1-Nrf2 pathway (Fan et al., 2010; Lau et al., 2010; Jain et al., 2010). These functions are dependent on p62’s unique ability to form dynamic cellular structures known as p62 bodies, through liquid-liquid phase separation (LLPS) (Sun et al., 2018).

p62 bodies are non-membrane-bound compartments that form through self-oligomerization mediated by its N-terminal PB1 domain (Ciuffa et al., 2015) and are further enhanced by the binding of p62 to the ubiquitin chains of ubiquitinated proteins via its C-terminal UBA domain (Zaffagnini et al., 2018). p62 bodies serve as essential platforms for both autophagosome formation and sequestration of Keap1, a key factor in antioxidant response pathways (Kageyama et al., 2021). The dynamic properties of p62 bodies are crucial determinants of their cellular functions. Recent studies have revealed that these structures are not static aggregates but undergo continuous fusion and reorganization (Zaffagnini et al., 2018; Sun et al., 2018), with their formation regulated by several factors. Negative regulators of p62 body formation include MOAP-1, a protein that binds to the PB1-ZZ domain of p62 (Tan et al., 2021); vault RNA1-1, a small non-coding RNA that interferes with p62 oligomerization (Horos et al., 2019); SPOP, an E3 ligase that ubiquitinates the UBA domain of p62 and decreases ubiquitin-binding capacity (Shi et al., 2022); and TRIM21, an E3 ligase that ubiquitinates the PB1 domain of p62 and inhibits oligomerization (Pan et al., 2016). Whereas several such factors negatively regulating p62 body assembly have been identified, the molecular mechanisms controlling the size and liquidity of p62 bodies and, consequently, the impact on Keap1 sequestration and downstream signaling functions, remain poorly understood.

Here, we identify SH3KBP1 binding protein 1 (SHKBP1) as a regulator of p62 body dynamics and function. We find that SHKBP1 directly binds to p62, and SHKBP1 levels inversely correlate with p62 body size and liquidity. Though SHKBP1 contains a Broad-Complex, Tramtrack and Bric a brac (BTB) domain and is thus a putative substrate adaptor of the Cullin-3 (CUL3) E3 ubiquitin ligase, we find that SHKBP1 does not affect p62 ubiquitination levels (Wang et al., 2020). Instead, loss-of-function studies reveal that, through these effects on p62 body dynamics, SHKBP1 inhibits the sequestration of Keap1 into p62 bodies and its subsequent clearance. Thus, our studies reveal an unexpected role for an E3 ligase substrate adaptor independent of ubiquitination in controlling the phase separation of p62 and clearance of Keap1.

## RESULTS

### SHKBP1 interactome analysis using quantitative proteomics identifies p62 as a potential binding partner

Our interest in SHKBP1 stemmed ultimately from a series of studies aimed to elucidate connections between phosphoinositide lipids and the regulation of the ubiquitination machinery. In this work, we focused on members of the PLEKHA family of pleckstrin homology domain-containing proteins, which through their multidomain interactions could link lipid binding to signaling via protein-protein interactions. Studies of PLEKHA4 revealed that it sequestered the substrate-specific adaptor of CUL3, KLHL12 in clusters at the plasma membrane via PI(4,5)P_2_ binding and oligomerization, leading to diminished polyubiquitination of Dishevelled proteins and enhancements in Wnt signaling and G1/S cell cycle progression in mammalian cells and *Drosophila* (Shami Shah et al., 2019, 2021; Sun et al., 2024). The paralogous PLEKHA5 interacted with and regulated a different Cullin family E3 ligase, the anaphase-promoting complex/cyclosome, albeit at microtubules, though the protein exhibited a cell cycle-dependent localization at the plasma membrane based on PI(4,5)P_2_ binding exclusively during mitosis (Cao et al., 2022b, 2024).

Analysis of the PLEKHA5 interactome by affinity purification–mass spectrometry proteomics revealed two additional high-confidence putative interaction partners, the paralogous SHKBP1 and KCTD3 (Cao et al., 2022b). Intriguingly, both of these proteins contain structural features that support a potential role as a CUL3 adaptor. Like KLHL12, they contain a BTB domain that binds to CUL3 (Pinkas et al., 2017; Balasco et al., 2024), and additionally they contain C-terminal WD40 repeats that likely mediate additional protein-protein interactions (Esposito et al., 2022, 2021; Ji et al., 2016; Pinkas et al., 2017). Mutations in SHKBP1 have been identified in multiple cancers (Cancer Genome Atlas Research Network et al., 2017; Angrisani et al., 2021), suggesting potentially important cellular roles. However, no SHKBP1- or KCTD3-dependent ubiquitination substrates, or other cellular or molecular functions, have been reported to date. Therefore, we began our studies by designing a series of proteomics studies aimed to identify putative ubiquitination substrates of SHKBP1 as a means to elucidate molecular and cellular functions of SHKBP1.

Due to the transient nature of E3–substrate interactions, identifying substrates of specific E3 ligases is a persistent challenge. We designed three distinct SILAC-based proteomics approaches aimed to stabilize interactions between adaptors and substrates either before or after ubiquitination events. First, we used MLN4924, a neddylation inhibitor that disrupts the NEDD8 conjugation pathway and causes accumulation of unstable Cullin-RING ligase (CRL) targets including but not limited to CUL3 (Tan et al., 2013). We perform such studies in SHKBP1 knockout (KO) cells generated using CRISPR/Cas9 mutagenesis engineered to stably express a GFP-tagged SHKBP1 for GFP-based affinity enrichment for mass spectrometry (MS) (Fig. 1A). Second, we used a polyubiquitin-binding domain as a “ligase trap” to increase affinity for ligase–substrate complexes (Watanabe et al., 2020) (Fig. 1B). By fusion of tandem ubiquitin-associated (UBA) domains from Rad23, which delivers ubiquitinated substrates to the 26S proteasome, to SHKBP1, we aimed to bias enrichment of SHKBP1 interactors toward those inherently transient ligase–substrate interactions (Fig. S1A, B). Third, we designed and used a point mutant of SHKBP1 incapable of binding to CUL3 to distinguish CUL3 complex components from substrates (Fig. 1C). The CUL3 binding-deficient mutant F44A was designed according to a reported point mutation in the BTB domain of KBTBD8 (Fig. S1C, D) (Werner et al., 2015).

**Figure 1.**
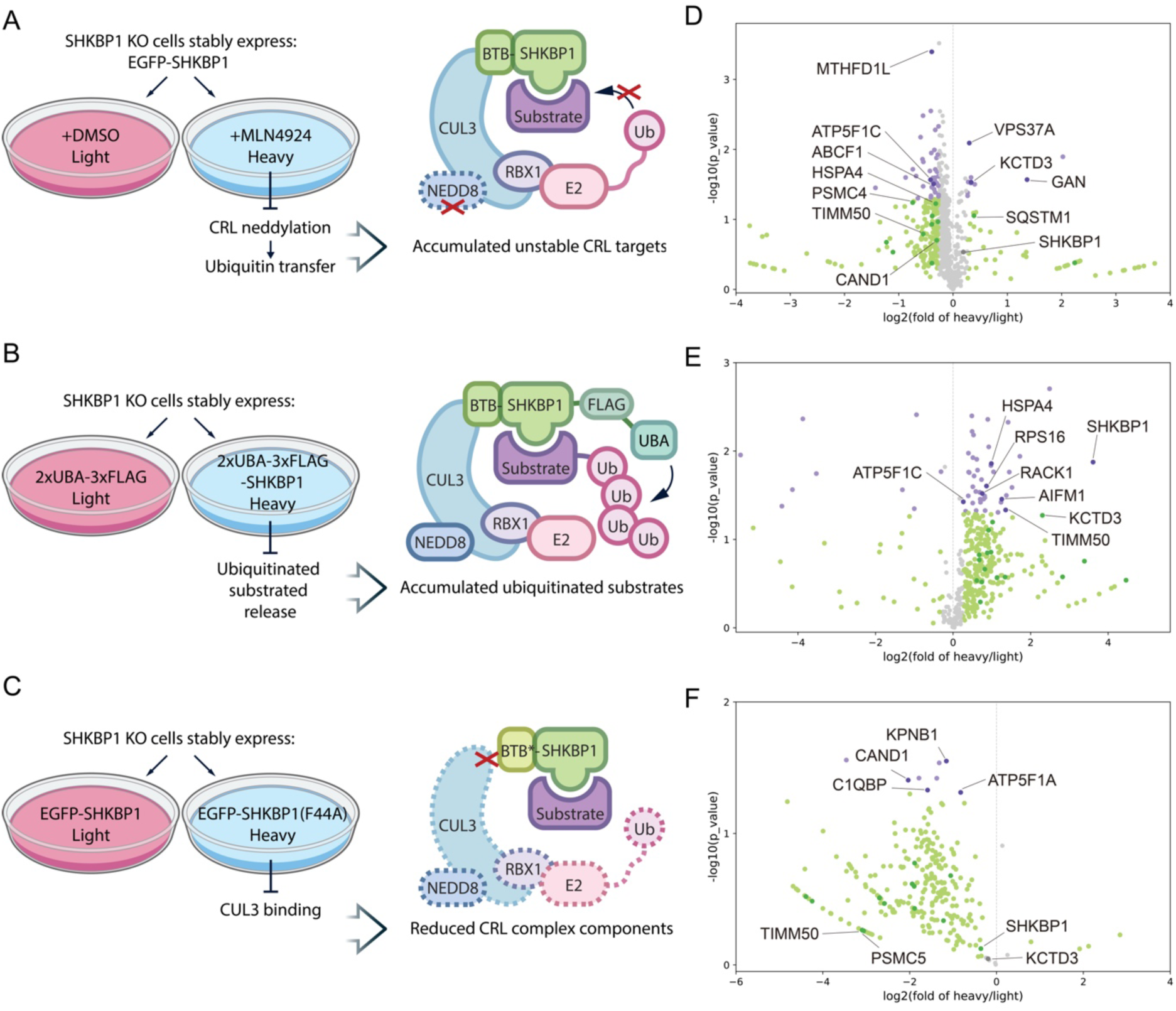
Quantitative affinity proteomics identifies SQSTM1/p62 as an interaction partner of the CUL3 adaptor SHKBP1. **(A)** Workflow of first proteomics experiments using the neddylation inhibitor MLN4924 to enrich unstable CRL targets (left) and cartoon representation of drug treatment mechanism (right). Cells were treated with MLN4924 (10 μM) or vehicle for 16 h. **(B)** Workflow of second proteomics experiments using tandem UBA domain fusions to SHKBP1 to enrich ubiquitinated substrates (left) and model of ligase trap (right). Cells stably expressing the corresponding construct were treated with MG-132 (20 μM) for 2 h. **(C)** Workflow of third proteomics experiments using the SHKBP1^F44A^ CUL3-binding deficient mutant to reduce CRL complex components in IP compared to SHKBP1^WT^ (left) and corresponding model (right). **(D–F)** Volcano plots from the three SILAC MS proteomics experiments, showing log_2_(fold changes of protein abundance in heavy/light samples) vs. statistical significance (–log_10_(p-value)). Proteins whose change was below the cutoff (fold change < 1.2) are indicated in grey. Those above the cutoff with p values above 0.05 are shown in green, and those with p values below 0.05 are shown in purple.

In all cases, we performed affinity enrichment of the tagged SHKBP1 proteins and, using SILAC-based quantitative proteomics, identified hits that would be predicted to be SHKBP1 interaction partners under the different conditions of each screen (i.e., + vs. – MLN4924, 2xUBA-SHKBP1 vs. 2xUBA only, and F44A vs. WT SHKBP1, respectively). From the combined results from these three sets of proteomics experiments, we identified several hits of interest (Fig. 1D–F). One group was additional CUL3 adaptors from the BTB family, including KCTD3, the closest paralog of SHKBP1, and GAN/KLHL16, which appeared as hits in the first two proteomics studies. Due to the propensity of BTB domains to homo- and hetero-oligomerize, it is possible that these BTB-containing proteins can hetero-oligomerize (Fig. S1E). A second group of hits were classified as CUL3 regulators. These include CAND1, a Cullin-RING exchange factor that preferentially binds to unneddylated cullins and, as expected, was more weakly enriched by SHKBP1^F44A^ compared to SHKBP1^WT^ in the third proteomics experiment, and the calcium sensor PDCD6, which has been implicated in the CUL3– KLHL12-mediated monoubiquitylation of SEC31 (Zheng et al., 2002; McGourty et al., 2016). A third and most diverse group was potential ubiquitination substrates, including SQSTM1/p62, TIMM50, ABCF1, AIFM1, VPS37A.

Among this third list, we were intrigued by the appearance of p62, which assembles into phase-separated p62 bodies to sequester intracellular polyubiquitinated cargos and mediate their autophagic clearance. Beyond its role in mediating autophagy of ubiquitinated proteins, p62 itself is heavily regulated by ubiquitination by several E3 ligases. At least five lysine residues within p62 (K7, K13, K91, K189, and K420) are targeted by at least four E3 ubiquitin ligases: TRIM21, Parkin, RNF166, and Keap1/SPOP, respectively (Pan et al., 2016; Song et al., 2016; Heath et al., 2016; Lee et al., 2017; Shi et al., 2022). We were therefore motivated to assess whether SHKBP1 could direct the ubiquitination of p62 by CUL3, acting as yet another layer of ubiquitin-related regulation of p62 function.

### SHKBP1 interacts with p62, and blocking neddylation increases this interaction and decreases p62 body size

We first validated the interaction between SHKBP1 and p62 through reciprocal co-immunoprecipitation (co-IP) experiments. In these experiments, forms of SHKBP1 and p62 tagged with either GFP or HA demonstrated mutual interaction (Fig. 2A, B). Given that p62 participates in degradation of ubiquitinated proteins and undergoes ubiquitination itself, we investigated the effects on this interaction by treatment of cells with either the proteasome inhibitor MG-132, to cause a buildup of ubiquitinated proteins destined for proteasomal degradation, and the neddylation inhibitor MLN4924, which prevents CRL-mediated ubiquitin ligase activity (Fig. 2C, D). Despite the differential changes in p62 protein levels caused by these treatments, they both resulted in robust SHKBP1 interactions with tagged p62 (Fig. 2C) and endogenous p62 (Fig. 2D).

**Figure 2.**
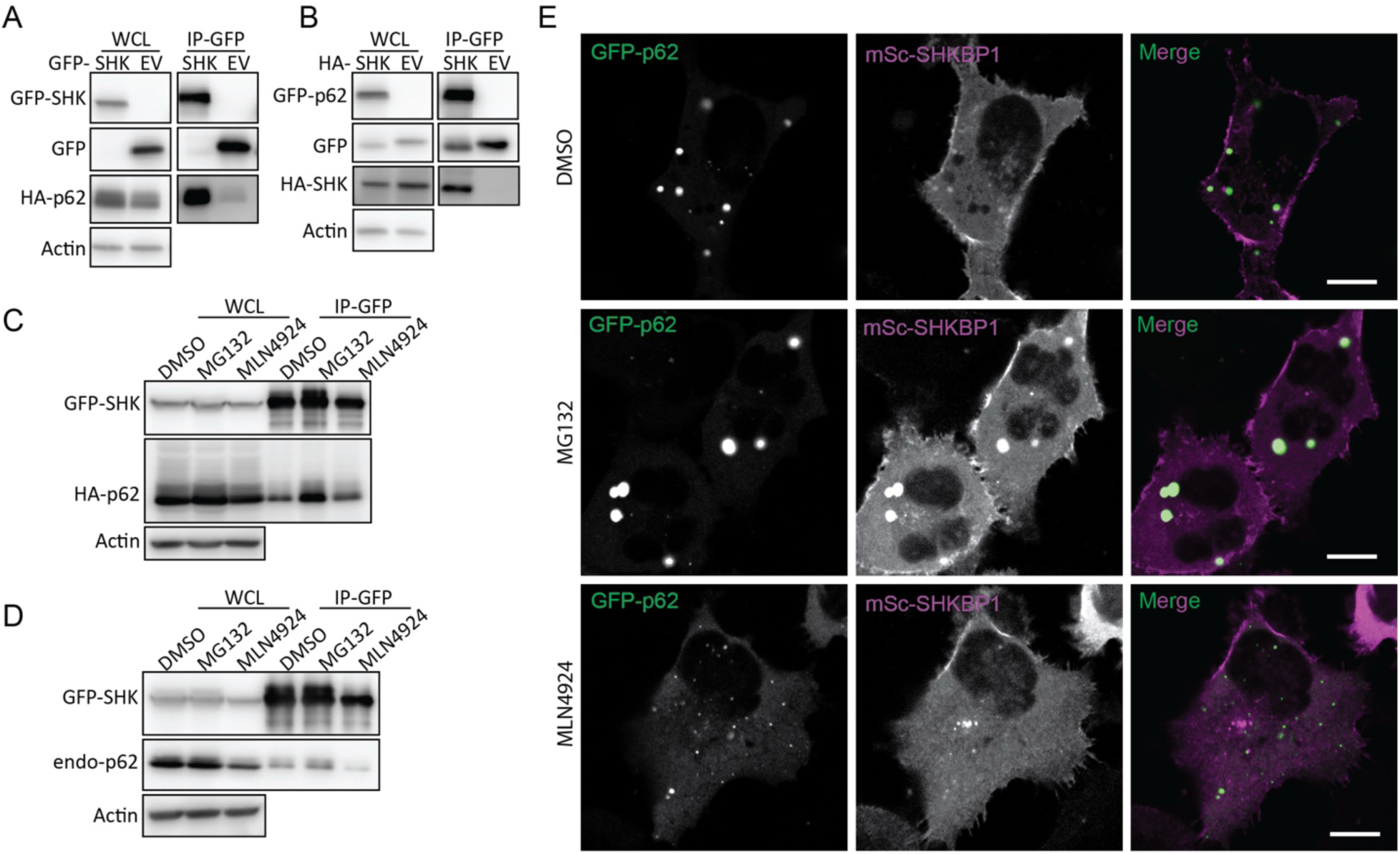
SHKBP1 interacts with p62, and this interaction is enhanced by MLN4924 treatment. **(A)** Western blot analysis of whole cell lysates (WCL) and α-GFP immunoprecipitates (IP) from HeLa cells co-transfected with HA-p62 and GFP-SHKBP1 or GFP empty vector (EV) as a control. **(B)** Western blot analysis of WCL and α-GFP IP from HeLa cells co-transfected with GFP-p62 and HA-SHKBP1 or HA empty vector (EV) as a control. **(C and D)** Western blot analysis of WCL and α-GFP IP to assess the interaction between SHKBP1 and exogenous (C) or endogenous p62 (D). HeLa cells were transfected with GFP-SHKBP1 in combination with HA-p62 (C) or alone (D), and treated with DMSO or the proteasome inhibitor MG-132 (20 μM) for 2 h or MLN4924 (10 μM) for 16 h. **(E)** HeLa cells were co-transfected with GFP-p62 and mScarlet-i-SHKBP1, treated with DMSO or the proteasome inhibitor MG-132 (20 μM) for 2 h or MLN4924 (10 μM) for 16 h, and then observed under confocal microscope 24 h post transfection. Scale bars: 10 μm.

Notably, these treatments influenced p62 body size. Confocal microscopy analysis of tagged SHKBP1 revealed localizations both at the plasma membrane (PM) and in the cytosol, with cytosolic puncta exhibiting partial colocalization with p62 bodies. MLN4924 treatment led to smaller p62 bodies, whereas MG-132 treatment caused larger p62 bodies to form (Fig. 2E). We reasoned that the decrease in global ubiquitination caused by MLN4924 would lead to reduced sequestration of ubiquitinated proteins in p62 bodies, whereas accumulation of ubiquitinated proteins caused by MG-132 treatment would necessitate larger p62 bodies for autophagy-mediated degradation. These results suggest that the interaction of p62 with SHKBP1 might be related to a varied extent of p62 body formation under stress.

### The WD domain of SHKBP1 interacts with the PB1 domain of p62

After confirming the SHKBP1–p62 interaction, we next mapped the domains in each protein responsible for mediating the interaction. SHKBP1 comprises an N-terminal BTB domain, a central WD domain with 8 repeats, and a C-terminal region. The BTB domain facilitates CUL3 binding, whereas the WD domains typically serves as a substrate interaction platform in other E3 ligases. To characterize SHKBP1 domain functions, we generated construct encoding either individual domains (BTB, WD, C-term) or single domain deletions (BTB+WD (BW) and WD+C terminus (WC)) (Fig. 3A). Interestingly, all six truncated constructs lost the PM localization characteristic of full-length SHKBP1, instead exhibiting distinct subcellular distribution patterns; BTB showed strong nuclear localization, BW formed many puncta, WD and WC formed fewer and smaller puncta, and C-term was cytosolic (Fig. 3B). We expressed these constructs in cells to assess their ability to co-immunoprecipitate with CUL3 and p62 (Fig. 3C, D). These studies indicated that BTB domain deletion substantially reduced CUL3 binding, and WD domain deletion significantly decreased p62 interaction. We note that due to the oligomerization property of both the PB1 domain of p62 and the BTB domain of SHKBP1, the excision of certain domains in truncation constructs, when overexpressed in cells containing low, endogenous levels of full-length, WT proteins, may not be sufficient to fully abrogate the ability of the overexpressed, tagged protein constructs to interact with one another. Analysis of colocalization of p62 with SHKBP1 truncations by confocal microscopy consistently supported these findings (Fig. S2A). Therefore, we conclude that the BTB domain of SHKBP1 mediates interaction with CUL3, and the WD domain of SHKBP1 interacts with p62.

**Figure 3.**
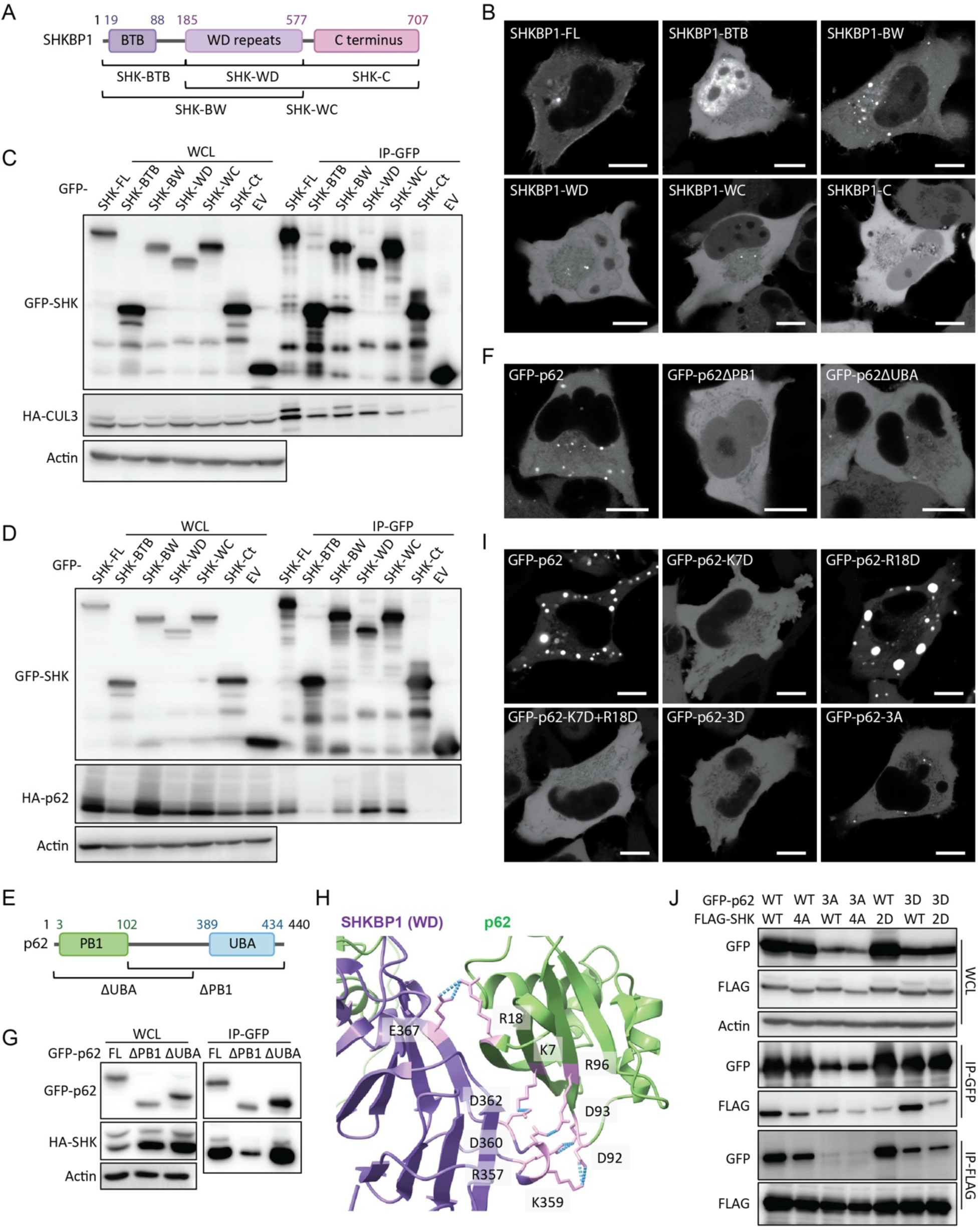
The WD domain of SHKBP1 interacts with the PB1 domain of p62. **(A)** Domain map of SHKBP1 and truncations used in this paper. **(B)** Representative live-cell images showing localization of full-length (FL) and truncated forms of SHKBP1. HeLa cells were transfected with the indicated GFP-tagged SHKBP1 construct and observed by confocal microscopy 24 h post-transfection. Scale bars: 10 μm. **(C)** Western blot analysis of α-GFP IP of lysates from HeLa cells co-transfected with HA-CUL3 and either the indicated GFP-SHKBP1 truncation or GFP empty vector (EV) as a control. **(D)** Western blot analysis of α-GFP IP of lysates from HeLa cells co-transfected with HA-p62 and either the indicated GFP-SHKBP1 truncation or GFP-EV as a control. **(E)** Domain map of p62 and truncations used in this paper. **(F)** Representative live cell images showing localization of p62 truncations. HeLa cells were transfected with the indicated GFP-tagged p62 construct and observed by confocal microscopy 24 h post-transfection. Scale bars: 10 μm. **(G)** Western blot analysis of α-GFP IP of lysates from HeLa cells co-transfected with HA-SHKBP1 and the indicated GFP-p62 construct. **(H)** AlphaFold3 structural prediction of the interaction between SHKBP1-WD domain (magenta) and p62 (green). Electrostatic interactions (blue) indicated between p62 residues K7, R18, D92, D93 and R96 and SHKBP1 residues R357, K359, D360, D362, and E367 (pink), respectively. ipTM = 0.73, pTM = 0.54. **(I)** Representative live cell images showing localization of p62 point mutants. HeLa cells were transfected with the indicated GFP-tagged p62 construct and observed by confocal microscopy 24 h post transfection. Scale bars: 10 μm. **(J)** Western blot analysis of α-GFP and α-FLAG IP of lysates from HeLa cells co-transfected with the indicated FLAG-SHKBP1 and GFP-p62 constructs.

Unlike SHKBP1, p62 has been well studied in terms of the functions of its individual domains. The two most important domains for p62 body formation are N-terminal Phox and Bem1 (PB1) domain and the C-terminal ubiquitin-associated (UBA) domain (Fig. 3E). The PB1 domain mediates self-oligomerization in a manner that is reduced by K7 ubiquitination (Wilson et al., 2003; Pan et al., 2016). The UBA domain mediates interactions with ubiquitinated substrates, and the binding affinity to ubiquitin chains is enhanced after S403 phosphorylation (Matsumoto et al., 2011). Both mechanisms contribute to p62 phase separation. Consequently, truncated versions of p62 lacking either of these domains could not form large p62 bodies, with ΔPB1 unable to form any p62 bodies at all and ΔUBA able to form fewer, and only smaller, p62 bodies (Fig. 3F). Co-IP experiments of these two truncations with SHKBP1 demonstrated that the deletion of the PB1 domain more strongly decreased the binding of p62 with SHKBP1 than deletion of the UBA domain, indicating that the PB1 domain is responsible for the interaction of p62 with SHKBP1 (Fig. 3G).

To further investigate the interaction interface between SHKBP1 and p62, we modeled the complex using AlphaFold 3 (Abramson et al., 2024), with full-length p62 (aa1-440) and the WD domain of SHKBP1 (aa185-577) used for the modeling. The predicted complex structure implicates an interface between the PB1 domain of p62 and the fourth WD repeat of SHKBP1 as primarily responsible for binding. Notably, five salt bridges were identified at the predicted interaction site (Fig. 3H). To experimentally test this model, we constructed point mutations targeting these salt bridge-forming residues. For p62, we generated a charge-eliminating mutant (D92A, D93A, R96A (3A)) and a charge-reversal mutant (K7D, R18D, R96D (3D)), and for SHKBP1, we generated similar charge-eliminating (R357A, K359A, D360A, D362A (4A)) and charge-reversal (R357D, K359D (2D)) mutants. In reciprocal co-IP experiments using GFP-tagged p62 and FLAG-tagged SHKBP1 constructs, we found that p62 mutations showed significantly lower protein levels (Fig. 3J). Comparison of the α-FLAG blots after α-GFP IP indicates that the SHKBP1-4A mutation significantly decreased the binding of SHKBP1 binding to p62 (lanes 1 and 2) and that binding affinity was further decreased between SHKBP1-4A and p62-3A (lanes 3 and 4). Similarly, the SHKBP1-2D mutation decreased the interaction between SHKBP1 and p62 (lanes 1 and 5) and even substantially with p62-3D (lanes 6 and 7).

Because p62 body formation appears to be related to the interaction with SHKBP1, we tested whether the p62 3A and 3D mutants also behaved abnormally in oligomerization using confocal microscopy analysis to assess subcellular localization. For these experiments, we not only used the 3A and 3D mutants of p62 but also K7D (which reverses charge at the known ubiquitination site), R18D, which was also predicted to interact with SHKBP1, and a K17D/R18D double mutant (Fig. 3I). Surprisingly, except for R18D, all mutations adopted a fully cytosolic localization, suggesting that these sites are important for the oligomerization function of the PB1 domain. Previous X-ray crystallography analysis of p62 has demonstrated that an intramolecular hydrogen bond between K7 and D69 is critical for p62 dimerization (Ichimura et al., 2008). Further, the E3 ligase TRIM21 modifies p62 at K7 with K63-linked polyubiquitin, which abrogates p62 oligomerization via the PB1 domain (Pan et al., 2016). Based on the above results, we hypothesize that the interaction with SHKBP1 via PB1 domain inhibits p62 oligomerization, thus negatively regulating p62 body formation.

### SHKBP1 inhibits p62 oligomerization without affecting its ubiquitination state

Given the potential of SHKBP1 to act as a CUL3-based E3 ligase substrate adaptor, we next examined whether p62 represents a CUL3–SHKBP1 ubiquitination substrate. We performed in vivo ubiquitination assays by transfecting cells with His-tagged ubiquitin and enriching ubiquitinated proteins using Ni-NTA affinity chromatography, with p62 ubiquitination detected as the ensemble of high-molecular weight species in the α-p62 blot (Fig. 4A). However, no significant differences in p62 ubiquitination were observed between WT cells, SHKBP1 KO cells, and WT cells overexpressing SHKBP1. We then explored the extent of p62 degradation and found that endogenous p62 protein levels decreased upon SHKBP1 KO and increased upon SHKBP1 overexpression (Fig. 4B). These results are consistent with a stabilizing physical interaction between these proteins, particularly through affecting the PB1 domain-mediated oligomerization mechanism.

**Figure 4.**
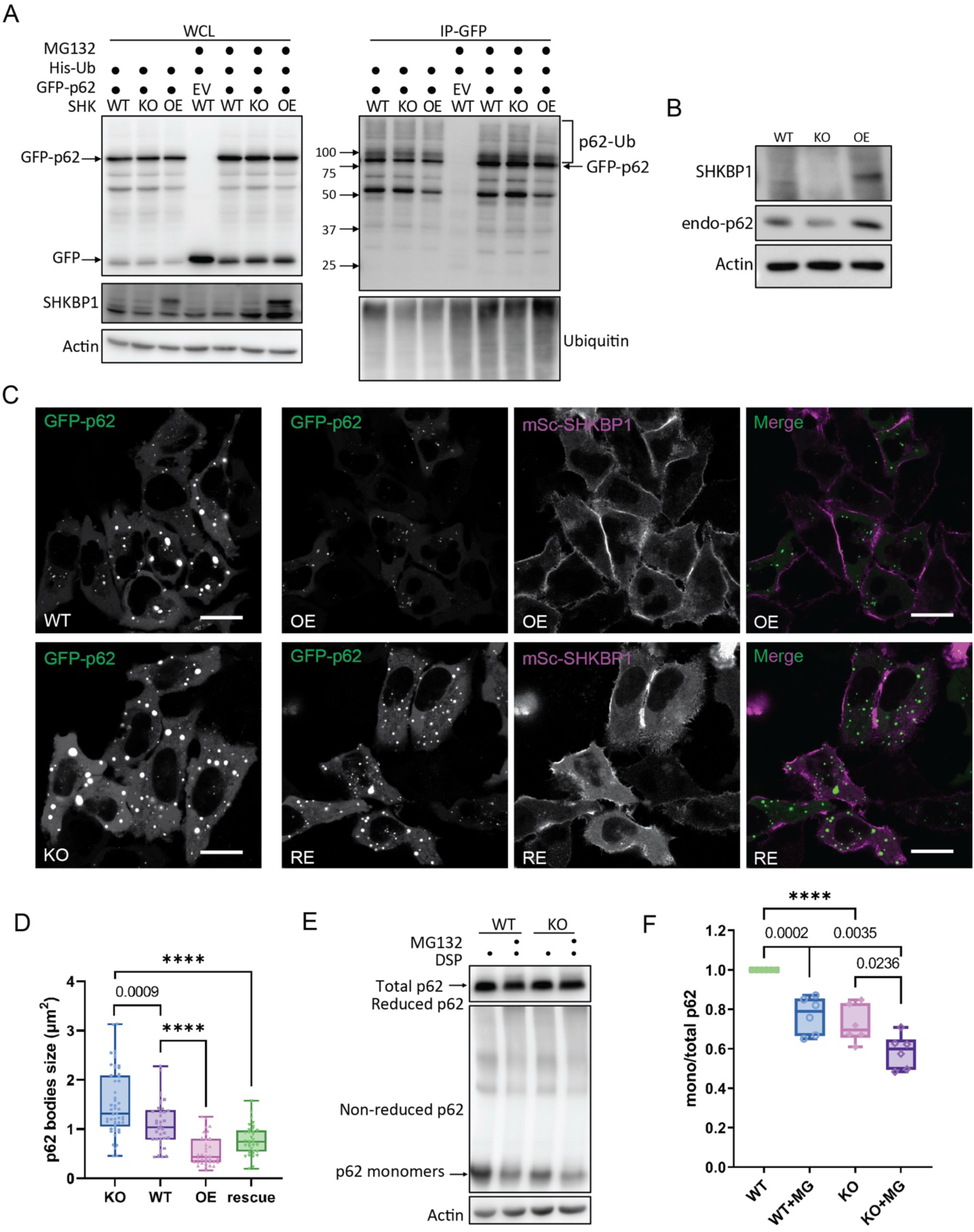
SHKBP1 inhibits p62 oligomerization without affecting its ubiquitination state. **(A)** Western blot analysis of in vivo ubiquitination assay. WT or SHKBP1 KO HeLa cells were co-transfected with His-ubiquitin and either GFP-p62 or GFP EV, and overexpressed (OE) group represents WT cells that were also co-transfected with FLAG-SHKBP1. 24 h post transfection, cells were treated with MG-132 (20 μM) for 2 h and subjected to His pull-down using Ni-NTA agarose and Western blot. **(B)** Western blot analysis of the whole-cell lysates (WCL) from WT, SHKBP1 KO, and FLAG-SHKBP1 overexpressing (OE) HeLa cells. **(C)** Representative live cell images showing p62 body formation. WT or SHKBP1 KO HeLa cells were transfected with GFP-p62 either alone (left) or in combination with mScarlet-i-SHKBP1 (right, where OE refers to SHKBP1 overexpression in WT cells and RE (rescue) refers to SHKBP1 overexpression in KO cells) and imaged using confocal microscopy 24 h post-transfection. Scale bars: 20 μm. **(D)** Quantification of the average p62 body size of images shown in (C) from three independently plated samples. n = 51 (KO), 43 (WT), 37 (OE), and 35 (RE). Exact p values indicated (****, p < 0.0001) from one-way ANOVA with Tukey’s post-hoc test. **E)** Western blot analysis of WCL from WT or SHKBP1 KO HeLa cells after DSP crosslinking. Cells were treated with MG-132 (0.5 μM) for 12 h, crosslinked with 0.4 mg/mL DSP at 4 °C for 2 h, and lysed in IP lysis buffer with 1% SDS. The lysates mixed with reducing or nonreducing loading buffer (i.e., with or without β-mercaptoethanol) and were analyzed by Western blot. **(F)** Quantification of the ratio of intensities of monomeric to total p62. Intensities were normalized to the WT without MG-132 treatment group. n = 6. Exact p values indicated (****, p < 0.0001) from two-way ANOVA with Tukey’s post-hoc test.

Oligomerization of p62 drives liquid–liquid phase separation (LLPS) to form p62 bodies, which have been considered as membraneless organelles (Komatsu, 2022). By confocal microscopy, we observed that SHKBP1 overexpression induced smaller p62 puncta compared to control cells, whereas conversely SHKBP1 KO cells displayed larger p62 puncta compared to WT cells in a manner that could be rescued by expression of a CRISPR-resistant SHKBP1 construct (Fig. 4C, D). We further examined p62 oligomerization biochemically, using the reversible crosslinking agent dithiobis(succinimidylpropionate) (DSP), which crosslinks lysine residues within or between proteins via a cleavable disulfide-containing linker. In the absence of reducing agents such as β-mercaptoethanol, large protein complexes migrate more slowly in SDS-PAGE upon DSP crosslinking, enabling the separation of monomers. To quantitatively assess p62 oligomerization, we treated WT and SHKBP1 KO cells with DSP and analyzed cell lysates under either reducing or nonreducing conditions. As a positive control, cells were also treated with MG-132 to induce proteotoxic stress, which as expected increased the extent of p62 oligomerization, which was quantified as a decrease in the ratio of p62 monomer to total p62 levels (Fig. 4E). We compared this ratio in SHKBP1 KO cells to that in WT cells and found that SHKBP1 KO led to a further decrease in p62 monomer levels and hence an increase in aggregation (Fig. 4F). Collectively, these data indicate that SHKBP1 inhibits p62 oligomerization in a ubiquitination-independent manner.

### SHKBP1 decreases p62 body liquidity

Biomolecular condensates, including p62 bodies, form through phase separation to create distinct cellular compartments that concentrate specific molecules (Shin and Brangwynne, 2017a). Previous studies have established that p62 forms droplets in vivo exhibiting liquid-like behavior that is characterized by high sphericity and the ability to undergo fusion and rapid molecular exchange with their surroundings (Sun et al., 2018).

To investigate how SHKBP1 influences p62 phase separation, we employed fluorescence recovery after photobleaching (FRAP) to quantify molecular dynamics within these cellular compartments. FRAP measures the rates of molecular exchange between p62 bodies and their surrounding environments, with faster rates of fluorescence recovery after local photobleaching indicating higher liquidity. To ensure consistent analysis across three different cell lines (WT, SHKBP1 KO, and SHKBP1-overexpressing HeLa cells), we examined spherical p62 bodies with approximately 2 µm diameter in all samples, as size determines surface area and can affect exchange rate and other dynamic properties. These studies revealed an inverse correlation between recovery kinetics and SHKBP1 expression status (Fig. 5A). p62 bodies in SHKBP1-overexpressing cells exhibited significantly slower fluorescence recovery rates compared to those in untransfected cells, whereas those in SHKBP1 KO cells exhibited accelerated recovery rates relative to those in WT cells. Further, p62 bodies in SHKBP1 KO cells had an increased mobile fraction of p62 relative to those in WT cells, whereas SHKBP1 overexpression increased the immobile fraction relative to untransfected cells (Fig. 5B). Recovery half-time comparisons further supported these observations (Fig. 5C). SHKBP1 KO cells showed significantly shorter half-times than WT cells and SHKBP1 overexpression. This enhanced molecular dynamics in KO cells aligned with our observation of an overall increase in p62 body size, as increased liquidity can promote condensate growth through more frequent molecular exchange and inter-condensate fusion (Xia et al., 2023; Shin and Brangwynne, 2017b).

**Figure 5.**
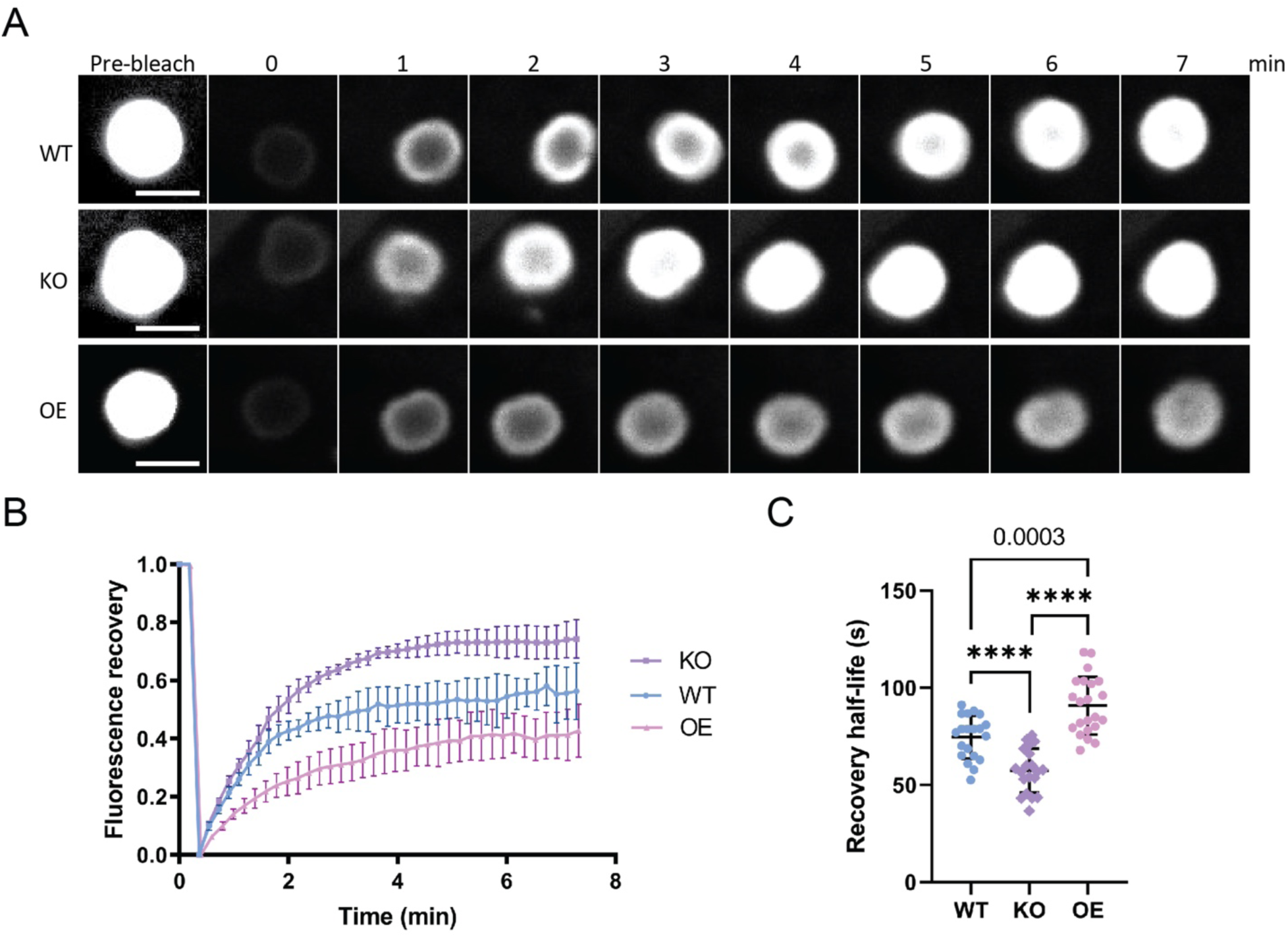
SHKBP1 decreases p62 body liquidity. **(A)** WT and SHKBP1 KO HeLa cells were transfected with GFP-p62 alone or in combination with mScarlet-i-SHKBP1. 24 h later, cells were subjected to live-cell imaging by confocal microscopy and analysis by fluorescence recovery after photobleaching (FRAP). Shown is a representative image for each condition. A pre-bleach image is provided along with a timecourse of post-bleach images (post-bleach time indicated in min). Scale bars: 2 μm. **(B)** Quantification of the fluorescence recovery rates of GFP-p62 from FRAP experiments. Data were analyzed with GraphPad Prism using nonlinear regression (curve fit), shown as mean ± standard deviation (SD). n = 3 for each group from three independent experiments. **(C)** Quantification of the half-life of fluorescence recovery of GFP-p62 from FRAP experiments. n = 20 for each group from three independent experiments. Exact p values indicated (****, p < 0.0001) from one-way ANOVA with Tukey’s post-hoc test.

These results suggest that SHKBP1 regulates p62 body liquidity by reducing molecular exchange rates. We propose that SHKBP1 achieves this negative regulation by modulating the dynamic exchange between two different pools of p62, namely the phase-separated p62 inside of p62 bodies and the pool of free p62 in the cytosol. In this model, SHKBP1 preferentially binds to monomeric, dimeric or low-complexity oligomeric p62, thereby inhibiting its further multimerization and ultimate incorporation into p62 bodies. This mechanism provides insight into how cells can tune the liquidity of p62 bodies through protein-protein interactions.

### SHKBP1 inhibits Keap1 aggregation and clearance via p62 bodies

p62 bodies play dual roles in cellular homeostasis by facilitating autophagic degradation and mediating antioxidant responses through the p62-Keap1-Nrf2 pathway. In this latter pathway, p62 competitively binds the CUL3 E3 ubiquitin ligase adaptor Keap1, preventing it from targeting the transcription factor Nrf2 for degradation, ultimately leading to Nrf2 nuclear translocation and activation of antioxidant gene expression (Komatsu, 2022; Baird and Yamamoto, 2020; Lau et al., 2010).

To investigate whether the interaction of SHKBP1 with p62 affects this antioxidant stress response, we first examined the binding between endogenous p62 and Keap1 in cells with different levels of SHKBP1. Enrichment of endogenous p62 by IP revealed that SHKBP1 KO enhanced p62-Keap1 binding relative to WT cells, whereas SHKBP1 overexpression reduced their interaction relative to untransfected cells, indicating that SHKBP1 antagonizes the association between endogenous p62 and Keap1 (Fig. 6A, B). We next explored the effect of SHKBP1 on Keap1 levels under conditions of oxidative stress by treatment of cells with sodium arsenite (As(III)) (Lau et al., 2013). Here, SHKBP1 KO cells exhibited enhanced Keap1 degradation both before and after As(III) treatment compared to WT cells (Fig. 6C, D).

**Figure 6.**
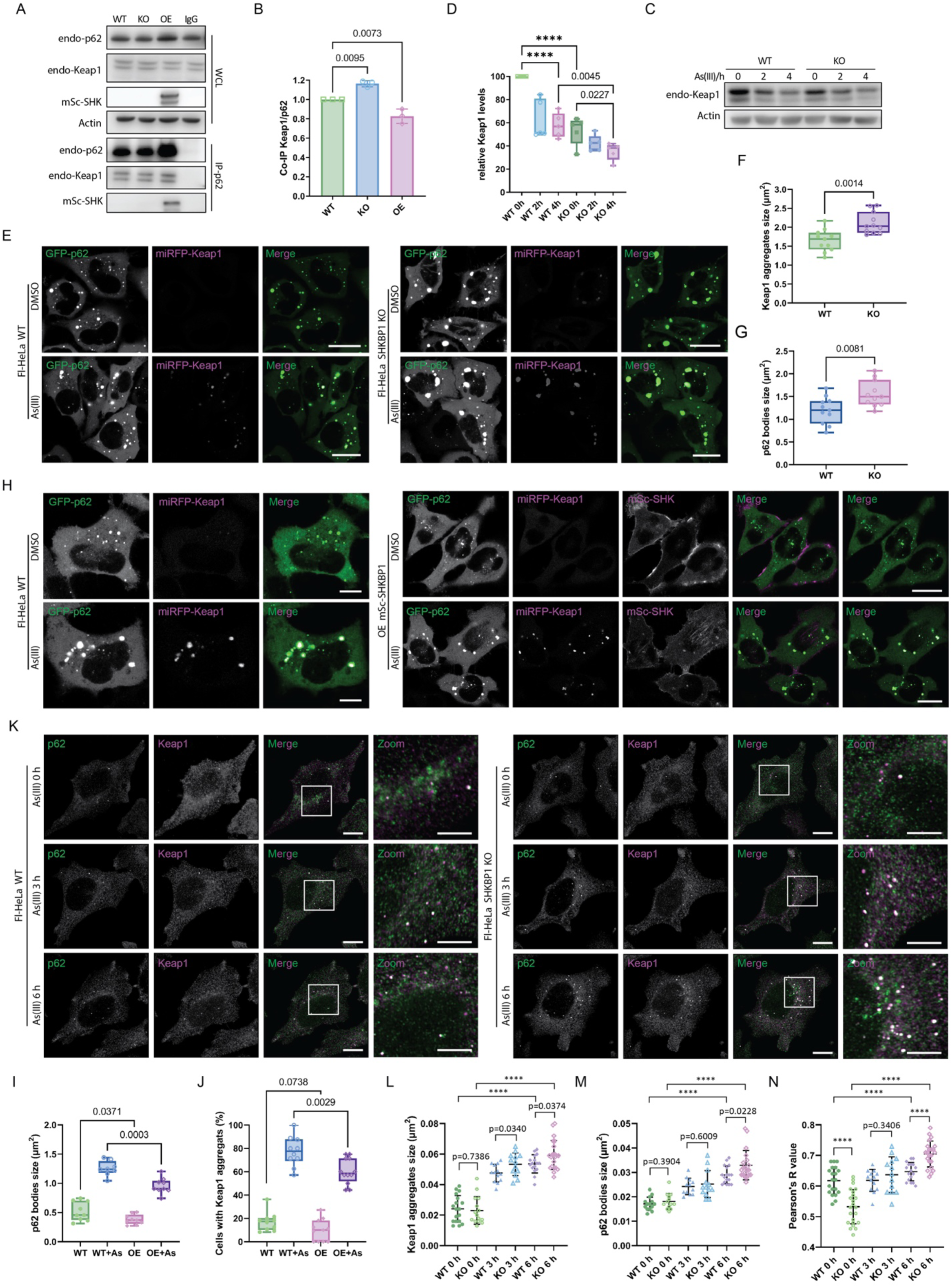
SHKBP1 inhibits Keap1 aggregation and clearance via p62 bodies. **(A)** Western blot analysis of co-IP experiments for endogenous p62. Lysates were from WT HeLa cells, SHKBP1 KO HeLa cells, and HeLa cells transfected with mScarlet-i-SHKBP1 (OE). **(B)** Quantification of Keap1 from p62-IP to demonstrate the interaction between endogenous p62 and Keap1, with band intensities normalized to actin levels (n = 3). Exact p values indicated from one-way ANOVA with Tukey’s post-hoc test. **C)** Western blot analysis of WCL from WT and SHKBP1 KO HeLa cells treated with As(III) (10 mM) for the indicated times. **(D)** Quantification of Keap1 protein levels, with band intensities normalized to actin levels (n = 5). Exact p values indicated (****, p < 0.0001) from two-way ANOVA with Tukey’s post-hoc test. **(E)** Confocal microscopy analysis of WT and KO SHKBP1 HeLa cells co-transfected with GFP-p62 and miRFP-Keap1 and treated with As(III) (10 mM) for 4 h or with DMSO as control. Scale bars: 20 μm. **(F and G)** Quantification of Keap1 aggregate size (F) and p62 body size (G) in WT and SHKBP1 KO HeLa cells after As(III) treatment (representative images shown in 6E). n = 11 images from three independently plated samples. Exact p values indicated from unpaired two-tailed Student’s t test with unequal variance. **(H)** Confocal microscopy analysis of HeLa cells co-transfected with GFP-p62 and miRFP-Keap1, with or without mScarlet-i-SHKBP1, and treated with As(III) (10 mM) for 4 h or with DMSO as control. Scale bars: 20 μm. **(I and J)** Quantification of p62 body size (I) and the percentage of cells containing Keap1 aggregates (J) in control of SHKBP1-overexpressing (OE) HeLa cells with or without As(III) treatment (representative images shown in 6H). n = 8-12 images from three independently plated samples. Exact p values indicated from two-way ANOVA with Tukey’s post-hoc test. **(K)** Immunofluorescence (IF) analysis of endogenous p62 and Keap1 in WT and SHKBP1 KO HeLa cells by confocal microscopy. Cells were treated with As(III) (10 mM) for indicated times. Scale bars: 10 μm; 5 μm (zoomed-in images). **(L-N)** Quantification of Keap1 aggregate size (L), p62 body size (M), and colocalization of Keap1 and p62 (N) in WT and SHKBP1 KO HeLa cells with As(III) treatment for the indicated times (representative images shown in 6K). n = 13-18 images from three independently plated samples. Exact p values indicated (****, p < 0.0001) from two-way ANOVA with Tukey’s post-hoc test.

Because oxidative stress induces Keap1 to form cytoplasmic aggregates that colocalize with p62 bodies before autophagic degradation, we quantified the size of both Keap1 aggregates and p62 bodies in cells co-expressing GFP-p62 and miRFP-Keap1 that contained different levels of SHKBP1. Compared to WT cells, SHKBP1 KO cells exhibited in larger Keap1 aggregates and p62 bodies (Fig. 6E–G), whereas SHKBP1 overexpression resulted in smaller p62 bodies and significantly reduced the percentage of cells containing Keap1 aggregates after As(III) treatment compared to untransfected cells (Fig. 6H–J). Consistently, a similar analysis of endogenous p62 and Keap1 by IF in SHKBP1 KO cells revealed larger p62 bodies and Keap1 aggregates than WT cells, accompanied by increased colocalization between Keap1 and p62 (Fig. 6K–N). Collectively, these findings demonstrate that SHKBP1 regulates p62-Keap1-Nrf2 signaling by modulating the ability of p62 to sequester Keap1, thereby influencing cellular antioxidant responses.

## DISCUSSION

p62 bodies are regulated in a multifaceted manner, including by phosphorylation, ubiquitination, and protein-protein interactions (Ichimura et al., 2013). Notable regulators including SPOP, TRIM21, MOAP-1, vault RNA1-1, DAXX, and Smurf1, all of which modulate p62 body formation through mechanistically distinct approaches (Shi et al., 2022; Pan et al., 2016; Tan et al., 2021; Horos et al., 2019; Yang et al., 2019; Xia et al., 2023). Compared to these established p62 regulators, SHKBP1 acts via a distinct and nuanced mechanism. Whereas it shares structural similarities with E3 ligase substrate receptors like SPOP and TRIM21, SHKBP1 does not mediate ubiquitination of p62 despite interacting with this protein. Instead, its interaction targets p62 oligomerization through direct interactions with relevant domains. The E3 ligase Smurf1 also interacts with p62 without ubiquitinating it, but its positive modulation on p62 body formation is indirectly achieved by an enhancement of p62 phosphorylation via mTORC1 signaling (Xia et al., 2023). MOAP-1 also binds to the PB1 domain, similar as SHKBP1, but it is recruited to p62 bodies to promote their dissociation upon stress (Tan et al., 2021). Instead, our data support a model wherein SHKBP1 interferes with p62 monomer oligomerization before condensate formation in a preventative mechanism that contrasts with direct engagement of MOAP-1 with p62 bodies.

Previous studies have highlighted that only monomeric and low-complexity oligomeric p62 demonstrate strong RNA binding (Horos et al., 2019), similar to the preference of SHKBP1 for binding to pools of p62 that are not phase-separated into p62 body. However, the cellular factors determining the selectivity of p62 for vtRNA1-1 remain unclear. By contrast, our data support a direct and structurally well-defined interaction of SHKBP1 with the PB1 domain of p62. Finally, the interactions of DAXX with p62 provide an interesting counterpoint to the proposed mechanism for SHKBP1 action. Whereas DAXX increases p62 body solidity by regulating p62 oligomerization and promoting p62 phase separation (Yang et al., 2019), SHKBP1 appears to constrain p62 body formation, as SHKBP1 KO cells exhibit increased material exchange of p62 into and out of liquid droplets. These seemingly opposite roles in liquid-liquid phase separation will inspire deeper investigation into the molecular mechanisms that collectively contribute to p62 phase separation. This intricate constellation of regulatory mechanisms indicates that p62 phase separation is a finely tuned process involving multiple molecular actors. Understanding the different contexts in which SHKBP1 contributes most strongly to this process could provide critical insights into cellular stress response and protein quality control mechanisms.

Beyond our primary findings, this study also reveals SHKBP1 as a BTB domain-containing protein with numerous molecular interactions. Our systematic proteomics experiments have identified multiple potential interactors beyond p62, suggesting interesting future directions. First, whereas we have confirmed binding of SHKBP1 to CUL3 (Pinkas et al., 2017; Esposito et al., 2022; Balasco et al., 2024), the functional assessment of the potential E3 ligase activity of CUL3–SHKBP1 remains unexplored. Proteins such as TIMM50, ABCF1, AIFM1, and VPS37A represent promising candidates to investigate potential ubiquitination mechanisms mediated by the CUL3–SHKBP1 complex. Second, the capacity of the BTB domain for multimerization presents additional avenues for investigation. BTB domain-mediated multimerization can significantly enhance local substrate concentration and ubiquitin transfer efficiency through homo- and hetero-multimerization mechanisms (Sumara et al., 2007; Frendo-Cumbo et al., 2019; Tong et al., 2006; Errington et al., 2012). Our identification of KCTD3 and GAN as SHKBP1 interactors is particularly noteworthy, given their shared structural characteristics: a BTB domain and a β-propeller architecture formed by either WD repeats (KCTD3) or Kelch repeats (GAN). These molecular features suggest the potential for SHKBP1 to form hetero-multimeric complexes, possibly while interacting with CUL3. Finally, mutations in SHKBP1 have been identified in cervical cancer (Greif et al., 2011), acute myeloid leukemia (Angrisani et al., 2021) and large intestine cancer (COSMIC database), suggesting potential roles of SHKBP1-mediated p62 dysfunction and/or the Keap1–Nrf2 antioxidant response pathway in tumorigenesis.

## MATERIALS AND METHODS

### Reagents

The following antibodies were used for Western blot: mouse anti-SQSTM1/p62 (D5L7G), Cell Signaling Technology (88588); rabbit anti SQSTM1/p62, ABclonal (A19700); mouse anti-Keap1, Santa Cruz Biotechnology (sc-365626); rabbit anti-SHKBP1, Invitrogen (PA521731); mouse anti-Ubiquitin, Santa Cruz Biotechnology (sc-8017); mouse anti-GFP, Santa Cruz Biotechnology (sc-9996); mouse anti-mCherry, Sigma-Aldrich (MAB131873); mouse anti-Actin, MP Bio (08691001); rabbit anti-FLAG, Sigma-Aldrich (F7425); mouse anti-HA, Covance (MMS-101R); The following antibodies were used for IF: rabbit anti-FLAG, Sigma-Aldrich (F7425); rabbit anti SQSTM1/p62, ABclonal (A19700); mouse anti-Keap1, Santa Cruz Biotechnology (sc-365626); goat anti-mouse HRP, Jackson ImmunoResearch Labs (115-035-146); goat anti-rabbit HRP, Jackson ImmunoResearch Labs (111-035-144).

Reagents were obtained from the following sources: DSP (Dithiobis(succinimidyl propionate)), TCI America (TCD2473); MLN4924, Cayman (15217); MG-132, Selleckchem (S2619); GFP-Trap magnetic agarose, ChromoTek (gtma-20); Protein G Sepharose, BioVision (6511); Ni-NTA agarose, QIAGEN (133203974); Anti-HA magnetic beads, Thermo Scientific (88836); Anti-FLAG magnetic beads, made in-house from NHS magnetic beads, Nacalai USA (TAS8848N1141-10MG) and Anti DYKDDDDK tag, Fujifilm (012-22383); Lipofectamine 2000, Invitrogen (11668019); RNase A, Research Products International (R21750); Puromycin dihydrochloride, Sigma-Aldrich (P8833); Blasticidin S hydrochloride, 10 mg/ml in HEPES buffer, Alfa Aesar (J67216); cOmplete Protease Inhibitor Cocktail, Roche (5056489001); ProLong Diamond Antifade Mountant with DAPI, Thermo Fisher (P36971); Clarity Western ECL Substrate, Bio-Rad (1705061).

### Cloning

The SHKBP1 cDNA (Gene ID: 92799, a gift from Haiyuan Yu, Cornell University) was cloned into the pEGFP-C1 vector (Clontech) using EcoRI and KpnI to generate EGFP-SHKBP1, and into the p3xFLAG-CMV-10 vector (Sigma-Aldrich) using HindIII and KpnI to generate FLAG-SHKBP1. Subsequently, truncations of SHKBP1 were generated by standard or overlap PCR, and inserts were cloned into the pEGFP-C1 and pmScarlet-i-C1 vectors, using EcoRI and KpnI. Mutant constructs were generated using the Quickchange XL site-directed mutagenesis kit (Agilent), including F44A, 4A, and 2D. For the ligase trap constructs, 2xUBA was subcloned from Rad23 into the p3xFLAG-CMV-14 vector (Sigma-Aldrich) and then SHKBP1 was inserted after the FLAG tag. For generating stable HeLa cells used in the proteomics experiments, silent mutations were incorporated into SHKBP1 to generate CRISPR KO SHKBP1 #1-resistant constructs using the Quickchange XL site-directed mutagenesis kit. Resistant EGFP-SHKBP1 or 2xUBA-3xFLAG-SHKBP1 (2US) or GFP-SHKBP1 (F44A) were subcloned into the pCDH-CMV-MCS vector to yield the lentiviral plasmid. p62-related constructs were amplified from HA-p62, Addgene (28027), which was subcloned into pEGFP-C1 vector (Clontech) using EcoRI and KpnI to generate GFP-p62. Subsequent truncations and mutations were generated similarly to those of SHKBP1.

### Cell culture and transfection

Flp-In T-Rex HeLa (Thermo Fisher) and HEK 293TN cells (Anthony Bretscher, Cornell University) were cultured in high-glucose DMEM (Corning) supplemented with 10% FBS (Corning) and 1% penicillin/streptomycin (P/S, Corning) at 37 °C in a 5% CO_2_ atmosphere. HEK 293TN cells were also supplemented with 1% sodium pyruvate (Corning). An SHKBP1 KO stable cell line was generated by CRISPR/Cas9-mediated mutagenesis in Flp-In T-Rex HeLa cells. Stable expression of GFP-SHKBP1, 2UF, 2US, and GFP-SHKBP1^F44A^ was achieved by lentiviral transduction as previously described (Cao et al., 2022a), followed by selection with puromycin (Sigma Aldrich) at 2 μg/mL or blasticidin (Alfa Aesar) at 50 μg/mL. Cell lines were used without further authentication, and mycoplasma testing (MycoSensor PCR assay, Agilent) was performed yearly. Plasmid transfections were performed using Lipofectamine 2000 (Invitrogen) following the manufacturer’s protocol. Cells were incubated with the transfection mix in Transfectagro (Corning) or Opti-MEM (Gibco) with 10% FBS for 6-8 h, and then the media was exchanged for regular growth medium.

### Immunoprecipitation and Western blots

For IP experiments, cells were harvested and lysed in IP lysis buffer (150 mM NaCl, 1% NP-40, 0.25% sodium deoxycholate, 5 mM EDTA, 50 mM Tris pH 7.5) supplemented with cOmplete protease inhibitor cocktail (Roche) on ice. Cell lysates were sonicated and centrifuged at 13,000 g for 5 min at 4 °C. Protein concentration of the clarified cell lysates was quantified using the BCA assay (Thermo Fisher). Approximately 10% of the clarified cell lysates was reserved, normalized, and used as input whole cell lysates (WCL). The rest was normalized and subjected to immunoprecipitation (IP) using anti-GFP or anti-FLAG or anti-HA beads with rotation at 4 °C overnight (12-16 h). The beads were washed three times with lysis buffer before denaturation at 95 °C for 5-10 min. WCL and IP samples were analyzed by SDS-PAGE and Western blot, with detection by chemiluminescence using Clarity Western ECL substrate (Bio-Rad), or by mass spectrometry-based proteomics analysis as described in more detail below. For IP of endogenous proteins, the clarified cell lysates were incubated with primary antibodies or IgG control for 4 h at 4 °C with rotation, followed by addition of Protein G Sepharose (BioVision) and rotation at 4 °C overnight. The resin was then centrifuged for 5 min at 1000 g, washed three times with lysis buffer, denatured and analyzed by SDS-PAGE and Western blot.

### SILAC labeling and mass spectrometry-based proteomics analysis

For SILAC proteomics, the indicated cell lines were cultured in heavy or light SILAC DMEM media (Thermo) supplemented with 10% dialyzed FBS (Corning) and 1% P/S for at least five passages to allow full labeling of cells before they were used in experiments. “Heavy” SILAC DMEM contained ^13^C_6_,^15^N_2_ L-lysine and ^13^C_6_,^15^N_4_ L-arginine. “Light” SILAC DMEM contained ^12^C_6_,^14^N_2_ L-lysine and ^12^C_6_,^14^N_4_ L-arginine. Samples for SILAC experiments were set up as follows: Experiment #1, GFP-SHKBP1^WT^ +MLN4924 (heavy) and GFP-SHKBP1^WT^ –MLN4924 (light); Experiment #2: 2xUBA-3xFLAG-SHKBP1 (heavy) and 2xUBA-3xFLAG (light); Experiment #3: GFP-SHKBP1^F44A^ (heavy) and GFP-SHKBP1^WT^ (light). Cells stably expressing the indicated construct were grown in the appropriate SILAC medium. For experiment #1, cells were incubated with MLN4924 (10 µM) or vehicle for 16 h. For experiment #2, cells were incubated with MG-132 (20 µM) for 2 h. Cells were collected, lysed in IP buffer, normalized, and then each pair of lysates from heavy and light cells within the same experiment were mixed together in a 1:1 ratio. Enrichment of GFP- or FLAG-tagged proteins was then performed by co-IP with GFP-Trap or anti-FLAG magnetic agarose as appropriate, as described above. Enriched proteins were eluted by denaturation at 95 °C in elution buffer (2% SDS (w/v), 20% glycerol (v/v), 300 mM Tris-HCl (pH 6.8)), followed by methanol/chloroform precipitation, reconstitution and disulfide reduction in 6 M urea, 10 mm DTT, and 50 mM Tris– HCl (pH 8.0), and alkylation with iodoacetamide (40 mM) for 1 h. Sample were diluted in 50 mM Tris–HCl (pH 8.0), trypsinized overnight, desalted using a Sep-Pak C18 cartridge, lyophilized, reconstituted, and analyzed by nanoLC-ESI-MS/MS on an UltiMate 3000 RSLCnano-Orbitrap Fusion system (Thermo Fisher Scientific). Each experiment was run in triplicate, with samples prepared on different days and analyzed by nanoLC-ES-MS/MS in series on the same day.

### Confocal microscopy

Cells were seeded on 35 mm glass-bottom culture dishes or on 12 mm cover glass in 12-well plates and then subjected to either live-cell imaging or fixation 24-30 h post-transfection for IF, respectively. Drug treatments including MG-132, MLN4924, and As(III), were performed right before harvest at the indicated concentrations and for the indicated times.

For IF analysis, cells were fixed in 4% paraformaldehyde in PBS for 15-20 min, followed by permeabilization and blocking with 0.1% Triton X-100 and 5% BSA in PBS for 30 min. Cells were subsequently incubated with primary antibodies diluted in blocking buffer for 1 h, then with Alexa Fluor-conjugated secondary antibodies diluted in blocking buffer for 1 h. All steps were performed at room temperature, and slides were stored at 4 °C for long-term storage prior to imaging.

Images were acquired via Zeiss Zen Blue 2.3 software on a Zeiss LSM 800 confocal laser scanning microscope equipped with 1.4 NA 40X and 1.4 NA 63X oil immersion objectives. Solid-state lasers (405, 488, 561, and 640 nm) were used to excite DAPI, EGFP/Alexa Fluor 488, mScarlet-i/Alexa Fluor 568, and miRFP respectively. Endogenous protein staining, including p62 and Keap1, were acquired using Airyscan.

### In vivo ubiquitination assays

Cells were transfected with His-Ubiquitin and other indicated constructs. 36 h after transfection, cells were harvested with PBS, 10% of cells were reserved for whole cell lysate (WCL), which was generated by lysis in RIPA lysis buffer (150 mM NaCl, 25 mM Tris pH 8.0, 1 mM EDTA, 1% Triton X-100, 0.5% sodium deoxycholate, and 0.1% SDS) supplemented with cOmplete protease inhibitor cocktail (Roche). The remaining 90% of cells were lysed in Buffer A (6 M guanidine-HCl, 0.1 M Na_2_HPO_4_/NaH_2_PO_4_, 10 mM imidazole, pH 8.0) by sonication and then incubated with equilibrated Ni-NTA-agarose (Qiagen) for 3 h at room temperature with agitation. The beads were washed once with Buffer A, then twice with Buffer B (10 mM Tris-Cl, pH 8.0, 8 M urea, 0.1 M NaH_2_PO_4_) and three times with a 1:4 dilution of Buffer B with an aqueous imidazole solution to a final concentration of 25 mM imidazole. The precipitate was resuspended in SDS loading buffer containing 200 mM imidazole, boiled at 95 °C for 10 min, and analyzed by Western blot.

### DSP crosslinking

Cells with at ∼80% confluence were washed twice with ice-cold washing buffer (10 mM Na_2_HPO_4_, 1.8 mM KH_2_PO_4_, 137 mM NaCl, 2.7 mM KCl, 0.1 mM CaCl_2_, and 1 mM MgCl_2_), and incubated in washing buffer containing 0.4 mg/ml DSP at 4°C for 2 h. After that, lysed cells with IP lysis buffer (150 mM NaCl, 1% NP-40, 0.25% sodium deoxycholate, 5 mM EDTA, 50 mM Tris pH 7.5) supplemented with 1% SDS. Lysates were clarified by centrifugation after a 30-min incubation on ice and then protein concentration was quantified using the BCA assay. Normalized WCL was then mixed with reducing or nonreducing loading buffer (i.e., with or without β-mercaptoethanol), denatured at 95 °C for 5 min, and analyzed by Western blot.

### Fluorescence recovery after photobleaching (FRAP)

HeLa cells were transfected with GFP-p62 alone or in combination with mScarlet-i-SHKBP1. 24-30 h post transfection, cells were subjected to FRAP analysis, conducted on an LSM 800 (Zeiss) using the FRAP module of the ZEN software, with an 1.4 NA 40X oil immersion objective. Bleaching was focused on circular regions of interest (ROIs) encompassing entire individual GFP-p62 puncta, which were bleached for 8 times with an interval of 10 s using the 488 nm laser at 100% intensity. Recovery was recorded for 7 min using Definite Focus to maintain z-position. Fluorescence intensities were analyzed by FIJI using the FRAP profiler plugin.

### Data processing, calculations, and statistics

All Western blot images and confocal microscopy images shown in figures are representative images from experiments performed in at least three biological replicates on different days. For Western blot quantification, band intensities were measured using FIJI. Statistical significance was calculated using one-way ANOVA or two-way ANOVA with Tukey’s multiple comparison test in GraphPad Prism. For quantification of confocal microscopy images, fluorescence intensities and particles sizes were measured using FIJI. Statistical significance was calculated with an unpaired two-tailed Student’s t-test with unequal variance or one-way ANOVA or two-way ANOVA with Tukey’s multiple comparison test as indicated. In figures containing bar graphs, each dot represents an individual biological replicate, the bar height represents the mean, and error bars represent the standard deviation. In figures containing scatter plots, each dot represents an image, the middle black line indicates the mean, and the top and bottom line indicate the standard deviation. In figures containing box graphs, each dot represents an individual biological replicate or image.

## ACKNOWLEDGMENTS

This work was supported by the National Institutes of Health (R01GM143367). We thank Sheng Zhang, Qin Fu, and Elizabeth Anderson from the Cornell Proteomics and Metabolomics Facility for performing SILAC proteomics studies. We thank Shun Enomoto, Jingyi Hu, Yizhen Jin, Adnan Shami Shah, and Hongyan Sun for technical assistance, the Fromme lab for access to equipment, and members of the Baskin lab for helpful discussions.

## AUTHOR CONTRIBUTIONS

L.L. and X.C. performed SILAC proteomics experiments. L.L. performed all other experiments. L.L. X.C., and J.M.B. conceived of the project idea, designed experiments, and analyzed data. L.L. and J.M.B. wrote the manuscript, and L.L., X.C., and J.M.B. edited the manuscript.

## SUPPLEMENTARY DATA

Table S1. Full datasets from proteomics experiments.

---

**Figure S1.**
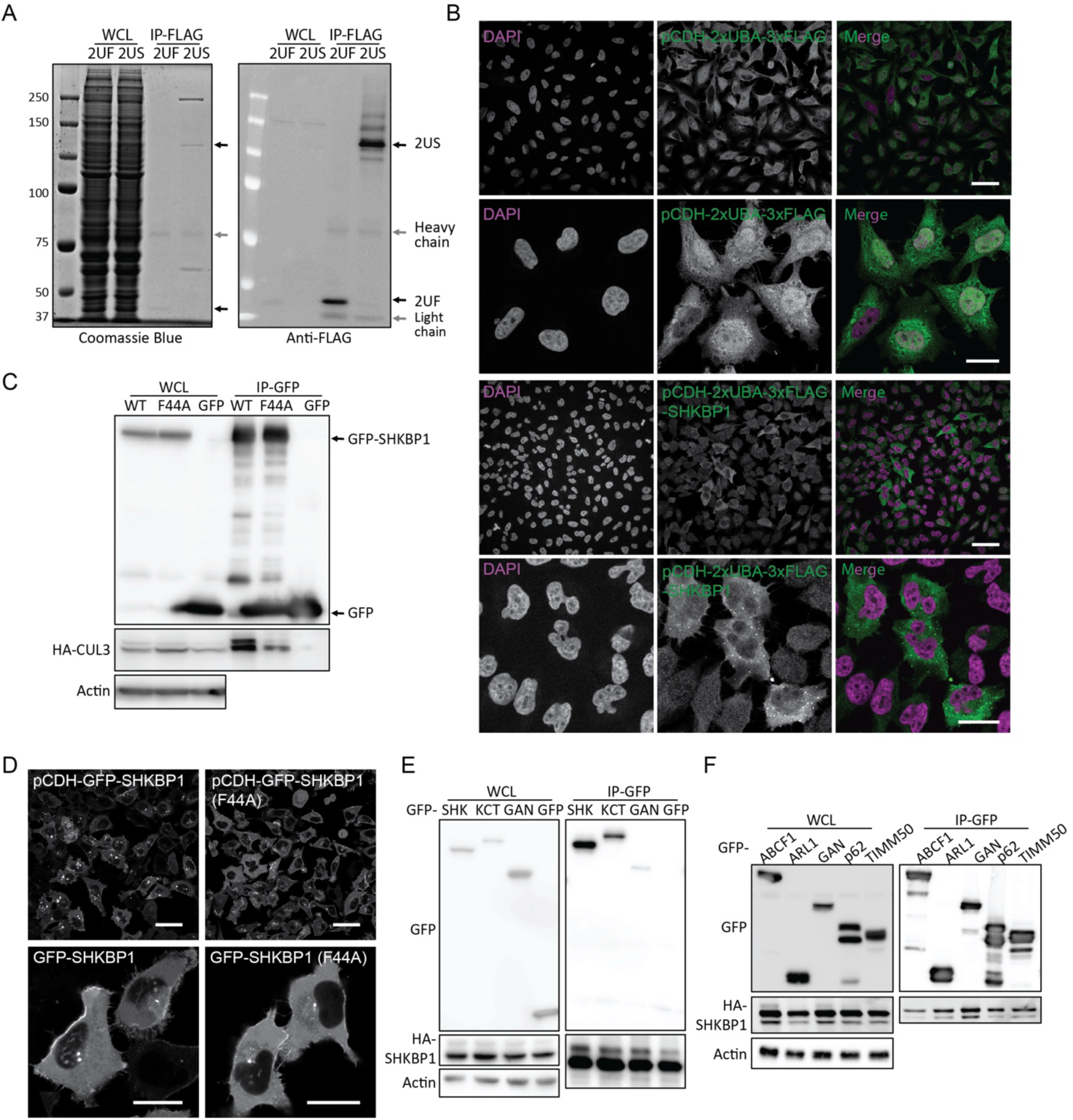
Preparation and verification of constructs and cell lines for proteomics studies. **(A)** Western blot analysis of whole cell lysates (WCL) and immunoprecipitates (IP) from Fl-HeLa cells stably expressing ligase trap constructs, showing proteins were significantly enriched in α-FLAG IP. 2UF, 2xUBA-3xFLAG; 2US, 2xUBA-3xFLAG-SHKBP1. **(B)** Immunofluorescence (IF) of cell lines stably expressing ligase trap constructs shown at low and high magnification. Cells were fixed, stained with α-FLAG (green) and DAPI (magenta), and imaged by confocal microscopy. Scale bars: 50 μm (first and third rows), 20 μm (second and fourth rows). **(C)** Western blot analysis of α-GFP IP from HeLa cells transfected with GFP only or GFP-tagged versions of SHKBP1^WT^ or SHKBP1^F44A^. **(D)** Confocal microscopy of HeLa cells stably expressing GFP-tagged SHKBP1^WT^ or SHKBP1^F44A^. Scale bars: 50 μm (top row), 20 μm (bottom row). **(E and F)** Western blot analysis of co-IP experiments for HA-SHKBP1 and GFP-tagged hits from proteomics to demonstrate the interaction between SHKBP1 and other BTB-containing proteins (E) and substrate candidates (F). SHK, SHKBP1; KCT, KCTD3.

**Figure S2.**
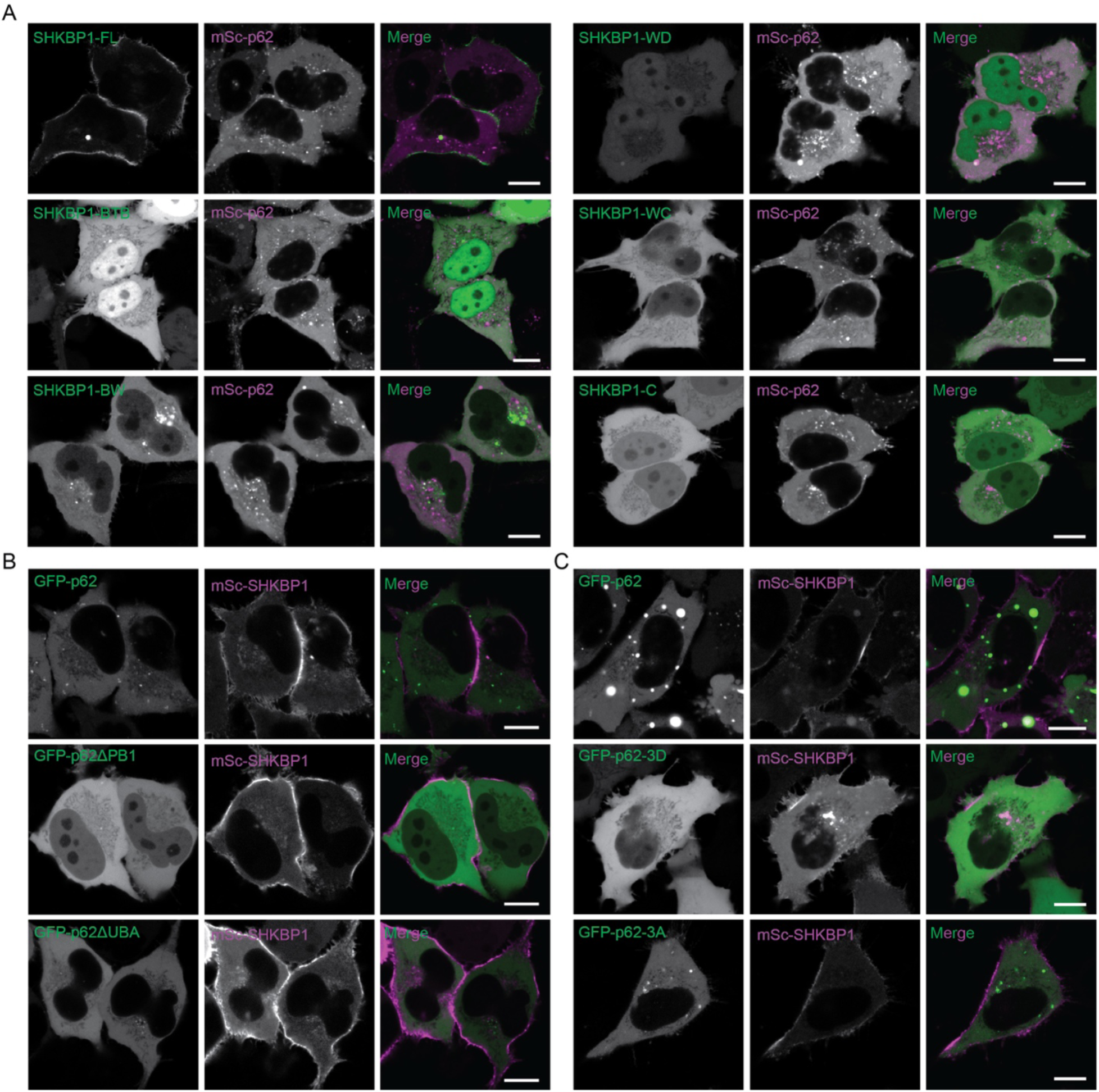
Intracellular localization of SHKBP1 and p62 truncations and mutants. **(A)** Confocal microscopy analysis of HeLa cells co-transfected with mScarlet-i-p62 (magenta) and the indicated GFP-SHKBP1 construct (green). **(B)** Confocal microscopy analysis of HeLa cells co-transfected with mScarlet-i-SHKBP1 (magenta) and the indicated GFP-p62 construct (green). **(C)** Confocal microscopy analysis of HeLa cells co-transfected with mScarlet-i-SHKBP1 (magenta) and the indicated GFP-p62 construct (green). Scale bars: 10 μm.

